# Cell Width in *Escherichia coli*: Interrogating Coupling to Nucleoid Structure

**DOI:** 10.1101/2023.05.15.540810

**Authors:** Charles B Howard, Avinoam Rabinovitch, Galit Yehezkel, Arieh Zaritsky

**Affiliations:** Department of Life Sciences, Faculty of Natural Sciences, Ben-Gurion University of the Negev, Be’er-Sheva 8410501 Israel; Department of Physics, Faculty of Natural Sciences, Ben-Gurion University of the Negev, Be’er-Sheva 8410501 Israel

**Keywords:** bacterial cell cycle / dimensions, division, growth, nucleoid complexity, replication positions

## Abstract

Cell dimensions of rod-shaped bacteria such as *Escherichia coli* are connected to mass growth and chromosome replication. It divides ≈20 min after termination of the replication cycle that initiates ≈40 min earlier at a relatively constant mass. Cells enlarge by elongation only, but at faster growth in richer media they are also wider. Width determination occurs in the divisome during the division process, coupled, temporally and spatially, to the ratio between the rates of growth and replication. The elusive signal directing the mechanism for width determination is related to the tightly linked duplications of the nucleoid (DNA) and the sacculus (peptidoglycan), the only two structures (macro-molecules) existing in a single copy. Six biologically meaningful parameters related to the key number of replication positions are reasonable candidates to convey such a signal. The current analysis discovered that of these, nucleoid complexity is the most likely parameter affecting cell width. As a corollary, a new, indirect approach to estimate replication rate is revealed.

---

Bacterial Physiology was established as a Quantitative Biology field in Copenhagen during the 1950’s and intensively overviewed then^1^ and again lately^2^. Its major outlook is the coordination between growth and the cell cycle in rigorous molecular biology and physics terms^3^. Elucidating the underlying principles of interactions between macromolecules, metabolites and structures clarifies some macroscopic and microscopic observations. Cell dimensions, however, are yet to be better explained, as described here.

Bacillary (rod-shaped) bacterial cells grow by elongation and divide perpendicularly at mid-point to two identical daughters. Under dynamic steady state^4^, cell mass (and volume) increases **exponentially** during its individual division cycle and in its culture mean size^5^—with doubling time (*τ* min) that depends on medium composition^6^. At 37°C, a cell divides ≈20 min (*D*) after termination of a round of replication that initiates ≈40 min (*C*) earlier at moderate to fast growth rates (*τ* < ≈70 min)^7^. Replication of the single, circular chromosome, bidirectionally from *oriC* to *terC*, is a **linear** process, initiating at a strain-specific constant cell mass per *oriC, Mi* or 2^n^-multiple [here, n=1,2,3, etc] of *Mi*^8^. Under conditions of *τ* < *C*, replication-initiation occurs before the previous replisome terminates its round, hence the multi-forked replicating chromosome contains more DNA. This model generates a larger mean cell *M* (= *Mi*×2^(*C*+*D*)/*τ*^) containing more DNA (in genome equivalents) *G* (= (*τ*/*C*ln2)×(2^(*C*+*D*)/*τ*^–2^*D*/*τ*^)) when growing faster in richer media^7^. Such cells are not only longer, by default, but surprisingly also wider^6,9^. The question whether the larger *G* in faster growing cells requires wider cells for proper segregation and partition before division is still moot. During nutritional shift-up transition^10^, cells overshoot their length during a ‘rate maintenance’ period of (*C*+*D*) before divisions are enhanced, and width *W* starts to expand at the divisome to form temporarily pear-shaped cells^11^. Thus, the tight spatial and temporal coupling between DNA replication / segregation and peptidoglycan synthesis is associated with determination of *W* during the division process itself.

There are six biologically meaningful parameters of chromosome content / structure known to us^12^,

i. the key number of “replication positions” *n* (= *C*/*τ*), and the other 5 that depend on it, as follows:
ii. the number of replicating “forks” *F* (= (2^*n*^–1); *i.e*., replisomes),
iii. the ratio between the frequency of *oriC* and that of *terC, o*/*t* (= 2^*n*^),
iv. the amount of DNA, in genome equivalents, associated with a single *terC* (*i.e*., a chromosome) *NC* (= (2^*n*^– 1) / (ln2×*n*))^13^, later termed “nucleoid complexity”^14^,
v. the cell’s “DNA concentration” *G*/*M* (= (1–2^-n^)/(*Mi*×*n*×ln2))^15^, and
vi. the amount of DNA per cell *G*^7^, the only one that depends on *D* as well.

Here, we attempt to find out which of these is the most likely parameter, statistically, that affects *W*. Table 1A summarizes (using Excel worksheets; see below and in Table S2 of the Supplementary material) the relationships between *W* and these 6 parameters in *E. coli* growing in the “mother machine” at 7 moderate and fast rates^16^, with *C*=40 min^7^. Exceedingly high linear correlation exists between *W* and *NC* (item iv above); the remarkably low coefficient of variation CV of 1.5% in the ratio *W*/*NC* (Table 1A) is rare in biological systems hence re-enforces the suggestion that it is not simply fortuitous. The outstandingly higher CV values (16.8%– 40.6%) of the ratios between *W* and each of the other 5 *n*-dependent parameters (items i-iii, v and vi) compellingly approves involvement of *NC* (iv) in *W* determination. In addition, *G*/*M* (v) can be rejected being inversely related to *W* (Table 1A).

**Table 1.** Correlations between cell width *W* and 6 *n*-dependent parameters related to nucleoid structure and replication assuming *C*=40 min

| **A (E.c.)** | $\tau$ | $W$ | $n$ | $F$ | $o/t$ | $NC$ | $G/M$ | $G$ | $W/n$ | $W/F$ | $W/(o/t)$ | $W/NC$ | $W/(G/M)$ | $W/G$ |
| --- | --- | --- | --- | --- | --- | --- | --- | --- | --- | --- | --- | --- | --- | --- |
|  | 17.40 | 0.98 | 2.299 | 3.921 | 4.921 | 2.460 | 0.500 | 5.458 | 0.426 | 0.250 | 0.199 | 0.398 | 1.960 | 0.180 |
|  | 22.22 | 0.80 | 1.800 | 2.483 | 3.483 | 1.990 | 0.571 | 3.713 | 0.444 | 0.322 | 0.230 | 0.402 | 1.400 | 0.215 |
|  | 26.67 | 0.72 | 1.500 | 1.828 | 2.828 | 1.758 | 0.622 | 2.957 | 0.480 | 0.394 | 0.255 | 0.409 | 1.158 | 0.243 |
|  | 30.77 | 0.67 | 1.300 | 1.462 | 2.462 | 1.623 | 0.659 | 2.546 | 0.515 | 0.458 | 0.272 | 0.413 | 1.017 | 0.263 |
|  | 37.50 | 0.61 | 1.067 | 1.095 | 2.095 | 1.480 | 0.707 | 2.143 | 0.572 | 0.557 | 0.291 | 0.412 | 0.863 | 0.285 |
|  | 50.85 | 0.55 | 0.787 | 0.725 | 1.725 | 1.330 | 0.771 | 1.746 | 0.699 | 0.759 | 0.319 | 0.414 | 0.714 | 0.315 |
|  | 51.28 | 0.55 | 0.780 | 0.717 | 1.717 | 1.326 | 0.772 | 1.738 | 0.705 | 0.767 | 0.320 | 0.415 | 0.712 | 0.316 |
| **Mean** |  |  |  |  |  |  |  |  | 0.549 | 0.501 | 0.269 | 0.409 | 1.118 | 0.260 |
| **SD** |  |  |  |  |  |  |  |  | 0.115 | 0.204 | 0.045 | 0.006 | 0.446 | 0.051 |
| **CV** |  |  |  |  |  |  |  |  | 0.209 | 0.406 | 0.168 | 0.015 | 0.399 | 0.196 |

| **B (S.t.)** | $\tau$ | $W$ | $n$ | $F$ | $o/t$ | $NC$ | $G/M$ | $G$ | $W/n$ | $W/F$ | $W/(o/t)$ | $W/NC$ | $W/(G/M)$ | $W/G$ |
| --- | --- | --- | --- | --- | --- | --- | --- | --- | --- | --- | --- | --- | --- | --- |
|  | 22.00 | 1.43 | 1.818 | 2.526 | 3.526 | 2.005 | 0.568 | 3.764 | 0.787 | 0.566 | 0.406 | 0.713 | 2.516 | 0.380 |
|  | 32.00 | 1.22 | 1.250 | 1.378 | 2.378 | 1.591 | 0.669 | 2.454 | 0.976 | 0.885 | 0.513 | 0.767 | 1.824 | 0.497 |
|  | 60.00 | 0.93 | 0.667 | 0.587 | 1.587 | 1.271 | 0.801 | 1.602 | 1.395 | 1.583 | 0.586 | 0.732 | 1.161 | 0.581 |
|  | 98.00 | 0.87 | 0.408 | 0.327 | 1.327 | 1.156 | 0.871 | 1.331 | 2.132 | 2.661 | 0.656 | 0.753 | 0.999 | 0.653 |
| **Mean** |  |  |  |  |  |  |  |  | 1.322 | 1.424 | 0.540 | 0.741 | 1.625 | 0.528 |
| **SD** |  |  |  |  |  |  |  |  | 0.596 | 0.928 | 0.107 | 0.024 | 0.693 | 0.117 |
| **CV** |  |  |  |  |  |  |  |  | 0.451 | 0.651 | 0.198 | 0.032 | 0.426 | 0.223 |

Values of cell width $W$ were measured in steady-state cultures of (A) *Eschericia coli*<sup>16</sup> and (B) *Salmonella typhimurium*<sup>6</sup> growing with the indicated doubling times $\tau$ (leftward block), calculated values of the 6 parameters (middle block, in green; see below and in text), and the respective values of the ratios between $W$ and each of the derived parameters (rightward block, in red) of the chromosome structure: $n$ (positions), $F$ (forks), $o/t$ (*oriC/terC* marker frequencies), $NC$ (nucleoid complexity, highlighted grey), $G/M$ (DNA concentration) and $G$ (DNA/cell). The ratios between $W$ and each of the 6 $n$ -dependent parameters and their statistics (in blue) were calculated using conventional values<sup>7</sup> of $C$ (40 min) and $D$ (20 min, for $G$ only).

A similar analysis was performed with data from the closely related species *Salmonella typhimurium* (Table 1B)^6^. The slightly higher CV (3.2%) of *W*/*NC* in this species than in *E. coli* may stem from at least 2 reasons: a smaller number of points (4 rather than 7), and the degree of accuracy in the method of measuring *W*. As observed in E. *coli*, the much higher CV values (19.8%–65.1%) of these 5 ratios (Table 1B) firmly support the major conclusion of this report, that a tight linear correlation exists between *W* and *NC*, at least in the gram-negative species studied here, *E. coli* and *S. typhimurium*.

Accurate and reproducible measurements of *W* are hampered by various methods of cell preparations and optical constraints (e.g.^18^). Values of *C* have been estimated using many modes (e.g.^7,15-17^; some reviewed in^14^), only two of which are direct, *in vivo*^19,20^. The uncertainties in values of *C*, the amazingly low CV of *W*/*NC* (Table 1), and the small number of parameters (*C* and *τ* only) involved in calculating *NC*, led us, serendipitously so, to assess if it is possible to obtain *C* by finding the minimal CV value of *W*/*NC* from the available measured data sets (*τ, W*). If *W* depends linearly on *NC, W* = *b* × *NC*, and we examine the change of the CV of *b* (= *W*/*NC*) for all cultures as a function of *C*, we get the best constant value of *W*/*NC* for the lowest CV. This happens for *E. coli* at *C* = 37.7 and *W*/*NC* = 0.42 (Table S1A) and for *S. typhimurium* at *C*= 37.5 and *W*/*NC* = 0.76 (Table S1B). Figure 1 (and the Supplementary Fig. S1) display the anticipated outcomes: the lowest CV (of 0.5% and 3%, respectively) occurred in both *E. coli* and *S. typhimurium* at an almost identical *C* of ≈37.6 min, remarkably close to the 39 min found recently^21^ using the accurate qPCR method, thus backing up, yet again, the *NC* as the decisive parameter for cell width determination.

**Figure 1.**
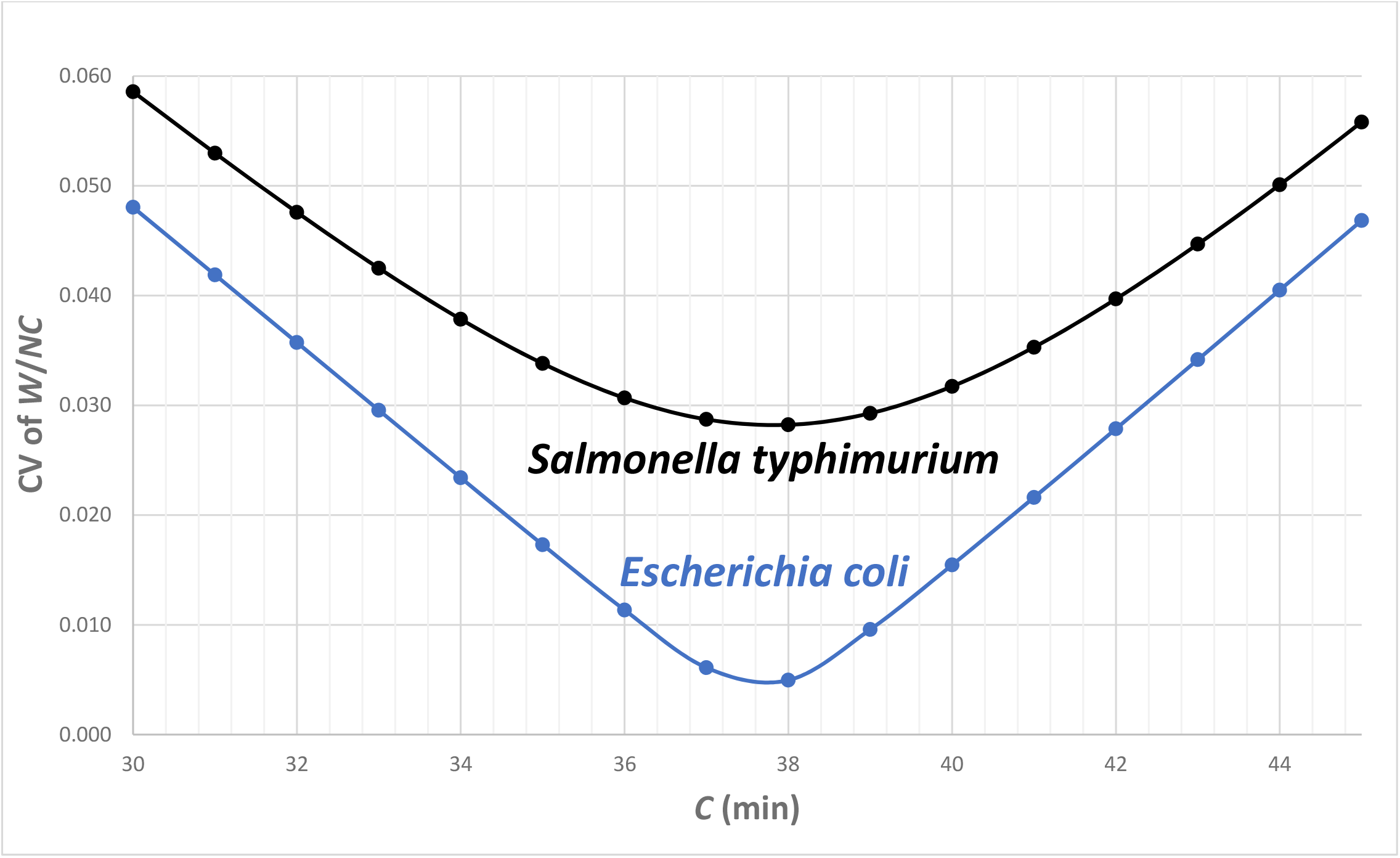
Values of CV of the ratio *W*/*NC* as a function of C extracted from Table S1

An additional set of data exists for another strain of *E. coli*, B/r H266 (Fig. 3C in^13^) rather than K12 NCM3722 of^16^, but the actual values of *W* in each of the 11 cultures (covering a wide range of *τ*’s) are not available (pers commun with CL Woldringh) and were cropped from the plot there^13^. A similar analysis was performed with these points (not shown), resulting in *C* = 47 min and CV of *W*/*NC* = 7.3%, substantially higher values than those obtained with the two major examples analyzed here (Table 1, Fig. 1 and Fig. S1), but considering the weak accuracy in the mode of cropping the measured *W* points, and the fact that they are still better off than all other *W*/parameter ratios, this result satisfactorily support our major conclusion about the relationship between *NC* and *W*. It is noteworthy that changes with *τ* in *W* (and even *C*) may differ among strains of the same species, which requires a separate analysis for each.

Coupling *W* to *NC via* the number of replication positions *n* (=*C*/*τ*) is supported also, at least qualitatively, by the dramatic changes in *W*^22^ upon modulating chromosome replication rate (*C*^-1^) by thymine concentration [T] supplied to *thyA* auxotroph^15^ (i.e., thymine limitation) without changing mass growth rate (60/*τ*). Quantifying this relationship is impeded by the lack of real steady state: under thymine limitation at short *τ*’s (< ∼50 min), cell size *M* changes continuously for long periods, and at a rate that is inversely related to [T]^22^. This change occurs firstly by increasing *W* up to a limit, then by branching^23^. The *W*-limit is consistent with the concept of existing eclipse^24^: a minimal distance from *oriC* needed for a replisome to reach before the next replisome can move forward to replicate the chromosome away from the new, temporarily abandoned initiating *oriC* (and see^25,26^). The recent finding that the rate of *in vivo* replication is not constant along the chromosome but rather oscillates, and in a still enigmatic manner^20^, complicates matters further, more so in thymine-limited slow replication rates as needed for accurate extensions of the analyses such as performed here. The option that the presumed signal for *W* determination is delivered at a specific time during the cell cycle (e.g., upon termination of replication) rather than by the mean value of *NC* has not skipped our mind; it will need a modified mode of calculations to that used here. For such analyses, additional results to the meagre that exist currently^27^ are needed.

The biochemical mechanism governing *W* in the gram-positive rod *Bacillus subtilis* is operated by the opposing action of the two main cell wall synthetic systems^28^. Similar correlation between *W* and directional MreB filament density / curvature in *E. coli* Rod mutants suggests that this model may be generalized to species that elongate *via* the Rod complex^29^, but the presumed primary signal affecting *W* by changing growth rate through the medium composition, let alone by thymine limitation, is still elusive.

Finally, understanding the mechanism of the elusive signal *NC*→*W* may affect our insight about the cellular activities during the *D* period. We shall not be surprised if deciphering the mechanism will lead to designing a new family of anti-bacterial drugs.

## Acknowledgements

This investigation was supported by grants to AZ from the U.S.-Israel Binational Science Foundation (BSF, No. 2017004) and the ISF-NSFC joint research program (No. 3320/20), and facilities by BGU administration. Charles E. Helmstetter and Conrad L. Woldringh are gratefully acknowledged for decades of inspiring cooperation and productive remarks in composing this manuscript.

**Figure S1.**
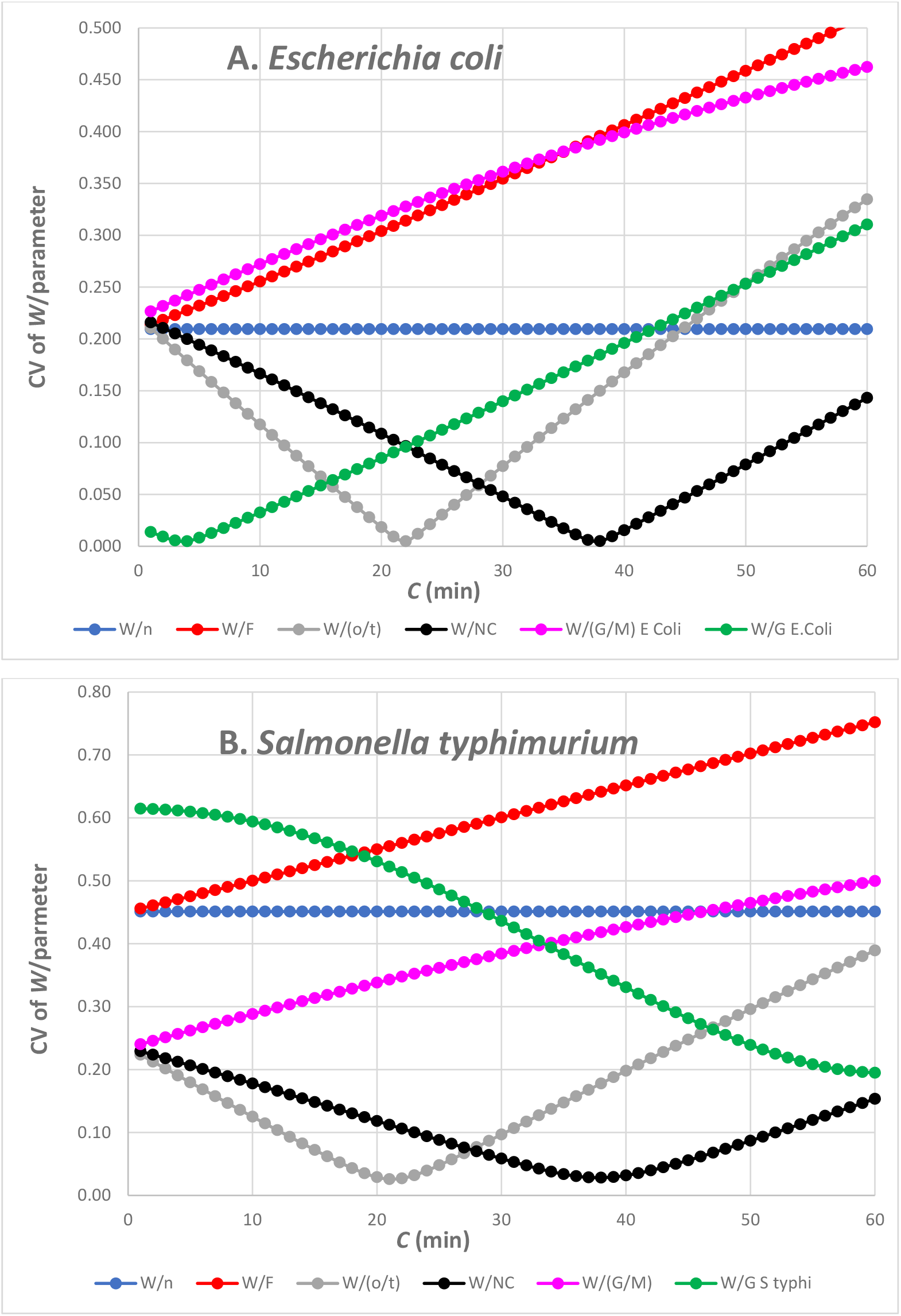
**A** and **B**: CV Values of each of the 6 *W*/parameter (i-vi) as a function of *C*, extracted from Table 1

## Supplementary Materials

**Figure S1, A** and **B**: above.

**Table S1**. Excel Worksheets: CV values of the ratios between *W* and each of the 6 parameters as a function of *C*, over the range 20 < *C* < 50 min, for *E. coli* (**A**) and *S. typhimurium* (B).

Table S1 (A and B) is in Excel format to ease analysis-usage by interested scientists. One must simply insert the values of two of the measured parameters of the three involved (*τ, W, C*) to obtain an approximate value for the third. Here, we used sets of known *τ* and *W* to estimate *C*, but the pairs (*τ, C*) would derive anticipated *W*.

